# SM3DD with Segmented PCA: A Comprehensive Method for Interpreting 3D Spatial Transcriptomics

**DOI:** 10.1101/2025.04.17.649456

**Authors:** Tony Blick, Aaron Kilgallon, James Monkman, Caroline Cooper, Chin Wee Tan, Emily E. Killingbeck, Liuliu Pan, Youngmi Kim, Yan Liang, Andy Nam, Michael Leon, Paulo Souza-Fonseca-Guimaraes, Seigo Nagashima, Ana Paula Camargo Martins, Cleber Machado-Souza, Lucia de Noronha, John F. Fraser, Gabrielle Belz, Fernando Souza-Fonseca-Guimaraes, Arutha Kulasinghe

**Author notes:** Corresponding author: Associate Professor Arutha Kulasinghe, Frazer Institute, Faculty of Health, Medicine and Behavioural Sciences, The University of Queensland, Translational Research Institute, 37 Kent Street, Woolloongabba, Queensland 4102, Australia.

## Abstract

We developed Standardised Minimum 3D Distance (SM3DD), an entirely cell segmentation/annotation-free approach to the analysis of spatial RNA datasets, using it to compare lung tissue from 16 clinically normal individuals to those of 18 SARS-CoV-2 patients who died from acute respiratory distress syndrome. RNA spatial coordinates were determined using the CosMx™ Spatial Molecular Imager (Bruker Spatial Biology, US). For each individual transcript location, we calculated the three-dimensional distances to the nearest transcript of each transcript type, standardising the distances to each transcript type. Mean SM3DDs were compared between normal and SARS-CoV-2 patients. Notably, hierarchical clustering of the directional log10(P) values organized genes by functionality, making it easier to interpret biological contexts and for FKBP11, where a decrease in distance to MZT2A was the most significant difference, suggesting a role in interferon signaling. Using a segmented principal components analysis of the entire SM3DD dataset, we identified multiple pathways, including ‘SARS-CoV-2 infection’, even though the assay did not include any SARS-CoV-2 transcripts.

## Introduction

Acute respiratory distress syndrome (ARDS) is, unsurprisingly, a serious complication of severe acute respiratory syndrome coronavirus 2 (SARS-CoV-2) infection. A global survey of ARDS incidence and outcomes in hospitalised SARS-CoV-2 patients found that, in a period prior to widespread vaccine use from the beginning of the pandemic through to the end of July 2020, approximately one third developed ARDS, and a mortality rate among ARDS-affected SARS-CoV-2 patients of 45% ^1^. Pulmonary damage from SARS-CoV-2 occurs through local and systemic inflammatory host responses, as well as through direct viral insult, with autopsy studies reporting alveolar damage and flooding, platelet microthrombi and extensive remodelling along with fibrotic evolution and variable collagen fibre deposition ^2^.

Spatially resolved transcriptomics, Method of the Year 2020, along with Spatial Proteomics, Method of the Year 2024, have transformed the analysis of tissue samples ^3, 4^. Currently, analysis methods for spatially resolved datasets predominantly rely on cell segmentation algorithms that are either geometry- or deep learning-based, but do not make use of transcriptomic data. These approaches rely heavily on nuclear detection with diminishing accuracy for tissue samples exhibiting a high degree of heterogeneity in cell sizes, shapes, orientations and densities within the extracellular matrix. Amongst other concerns, inaccurate cell segmentation can lead to mixed-cell expression profiles and subsequently errors in both cell type annotation and cell type-specific differential expression analysis.

There are currently few published methods for the analysis of large spatially resolved transcriptomic datasets that don’t rely on spatial restriction approaches to cell segmentation prior to the assignment of expression data to cells. Spot-based Spatial cell-type Analysis by Multidimensional mRNA density estimation (SSAM) instead assigns cell types at a pixel level based on a sliding window analysis of the spatially resolved transcriptomic data, resulting in improved cell type detection and assignment ^5^. Factor Inference of Cartographic Transcriptome at Ultra-high Resolution (FICTURE) also makes use of the spatial expression data to improve cell segmentation accuracy, deriving fine-scale tissue structure by identifying factors derived from gene expression patterns that fit to a Latent Dirichlet Allocation (LDA) model and adaptively aggregating pixel level information using anchors ^6^. The graph neural network model GraphSAGE ^7^ has been applied without cell segmentation to a pulmonary fibrosis spatial RNA dataset to describe molecular niches ^8^.

We present here Standardised Minimum 3D Distance (SM3DD), a novel cell segmentation/annotation-free analytical methodology for large spatially resolved transcriptomic datasets, which makes full use of the three-dimensional nature of the datasets, a benefit over current analysis approaches that flatten the data. Simplistically, the three-dimensional distance to the nearest transcript of each transcript type is calculated for every transcript location in the dataset. Within each field of view (FOV), the distances to each transcript type are standardized. For each sample, the mean SM3DD for each directional transcript type-to-transcript type pair is calculated, prior to a comparison between sample groups. Additionally, a principal components analysis (PCA) of the entirety of the SM3DDs is performed, segmenting the analysis according to the transcript type from which the 3D distances were determined, and mean differences in dimension scores across the PCA segments calculated, at which stage pathway level analysis can be applied. We used SM3DD to compare RNA spatial coordinates, generated using NanoString’s CosMx™ Spatial Molecular Imager 1000-plex assay ^9^, of lung tissue from 16 normal cases to that from 18 SARS-CoV-2 patients who died from ARDS. We further demonstrate the broad utility of SM3DD by applying it to a publicly-available Xenium 343-plex spatial RNA dataset for matched fibrotic regions from 8 patients with pulmonary fibrosis ^8^. We demonstrate that SM3DD facilitates detailed pathway level interpretation of spatial transcriptomic data.

## Results

### CosMx™ Spatial Molecular Image analysis of tissue microarrays

Tissue blocks of rapid autopsy lung samples from 18 SARS-CoV-2 patients, confirmed by RTqPCR of nasopharyngeal swabs, who died from ARDS were reviewed by an anatomical pathologist and 2 tissue microarrays (TMAs) constructed from 30 lung tissue cores. Autopsy samples were obtained from the Pontificia Universidade Catolica do Parana PUCPR the National Commission for Research Ethics (ethics protocol number 3.944.734/2020), with families permitted post-mortem biopsies and ratification by the University of Queensland Human Research Ethics Committee. A normal lung TMA (‘LCN241’, US Biomax, USA) was included as comparator samples. Adjacent serial tissue sections from all TMAs were profiled using the CosMx™ Spatial Molecular Imager (SMI) 1000-plex assay (Bruker Spatial Biology, USA). Fields of view (FOVs) were in matching locations across adjacent serial tissue sections, with data collected from a total of 116 FOVs from the SARS-CoV-2 patient lung samples and 38 FOVs across 16 independent normal lung samples. Four of the FOVs from one core were excluded from the analysis due to low transcript counts, resulting in a final total of 98,815,173 transcripts.

### Calculation of SM3DD

Within each FOV, for all transcript locations, the spatial coordinates, adjusted for Z scaling, were used to calculate the 3D distances to the nearest transcript of each transcript type, resulting in 96,838,869,540 directional transcript-to-transcript distances. These distances were standardised within each FOV, to correct for overall expression level differences and signal strength affects, by dividing the distances to each transcript type by the mean of the distances to that transcript type (Fig. 1).

**Figure 1.**
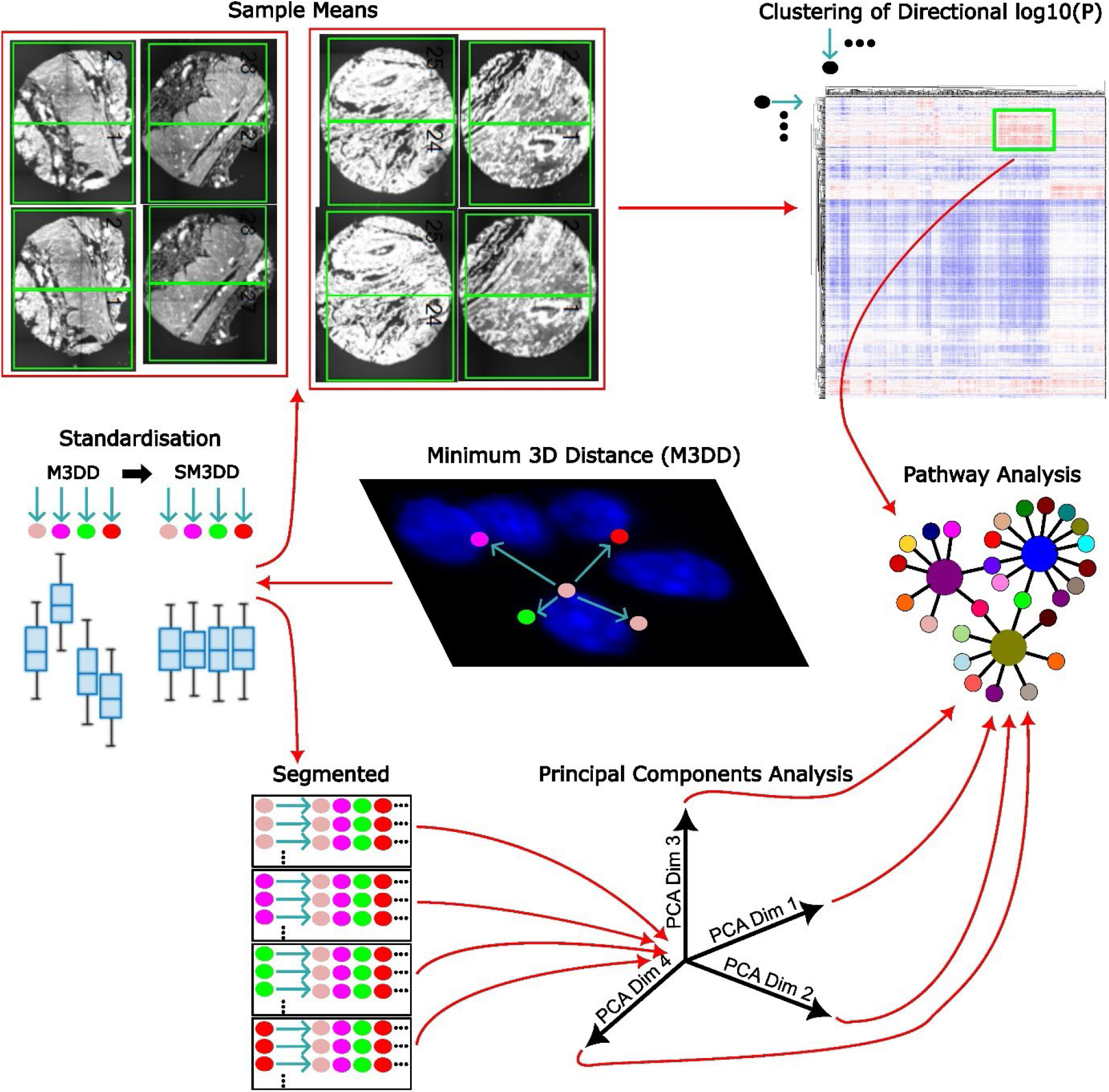
Method overview. The 3D distances to the nearest transcript of each transcript type (M3DD) are determined for all transcript locations. These distances are standardised (SM3DD) within each FOV by dividing by the mean of the distance to each transcript type. Sample means are determined and compared between groups by t-test and the directional log_10_(P) values clustered. Prominent clusters are assessed by Fisher’s Exact Test for pathway level effects. SM3DDs are also segmented according to the transcript type from which they were determined and PCA performed. Within each PCA dimension, differences in mean PCA scores between groups are calculated for each transcript, and these differences assessed by t-test for pathway level effects.

### Mean SM3DD comparison between SARS-CoV-2 and normal

FOVs were pooled and sample level means for each transcript type-to-transcript type SM3DD determined. Differences between mean SM3DDs for SARS-CoV-2 and normal samples were assessed by t-test, with the false discovery rate (FDR) controlled by the two-stage Benjamini, Krieger, and Yekutieli procedure ^10^, implemented by the MATLAB function fdr_bh ^11^. Approximately 45% (431,569) of all transcript type-to-transcript type mean SM3DDs differed between SARS-CoV-2 and normal samples for an FDR cut-off of 5% (Fig. 2A). Statistically, the most marked difference was the overall shorter FKBP prolyl isomerase (FKBP11)-to-mitotic spindle organizing protein 2A (MZT2A) transcript SM3DD in SARS-CoV-2 samples (P = 1.06 x 10^-17^). Across all mean SM3DDs specifically to MZT2A, this difference was very distinct (Suppl. Fig. 1). An overall shorter mean SM3DD to FKBP11 transcripts from nuclear paraspeckle assembly transcript 1 (NEAT1) was statistically ranked 13^th^ (P = 2.04 x 10^-13^), which is notable as NEAT1 is non-coding and nuclear localised.

**Figure 2.**
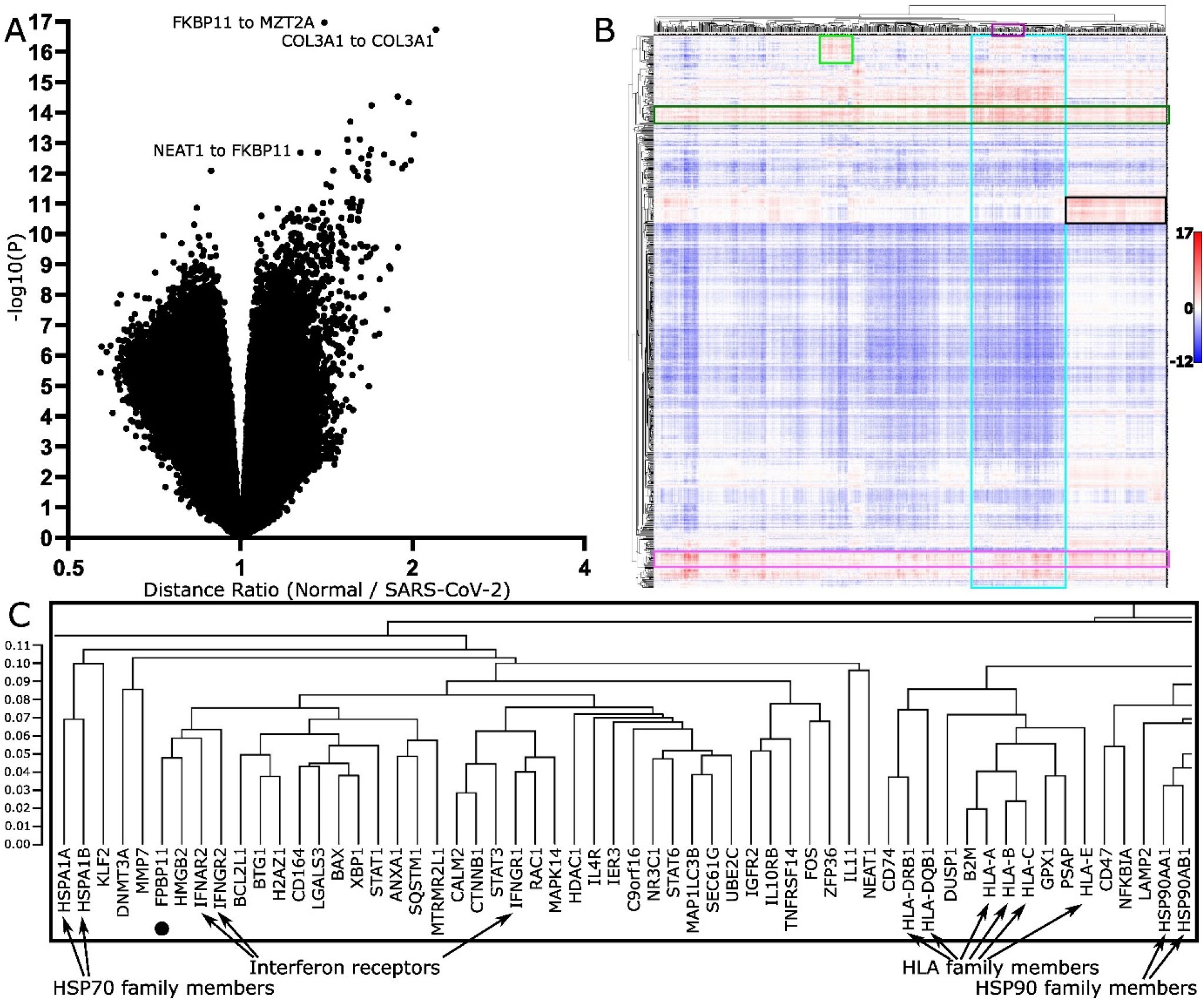
Comparison of transcript distances between normal and Sars-CoV-2 infected lungs. (A) Volcano plot of all directional transcript distance comparisons. X-axis is distance ratio, where larger values represent shorter distances in Sars-CoV-2 infected lungs. Y-axes are negative log10 (P values). (B) Clustering of directional negative log10(P values) from comparison of transcript distances between normal and Sars-CoV-2 infected lungs. Scalebar indicates directional negative log10(P) value. Prominent clusters of transcripts with shorter distances (red) in Sars-CoV-2 infected lungs were assessed by pathway analysis, identifying that these likely represent widespread interferon γ signaling (dark green box) and extracellular matrix (ECM) deposition (pink box), along with pathogen phagocytosis (light green box), ‘SARS-CoV-2 infection’ and an extensive list of predominantly signaling pathways (blue box) and, although not statistically significant, what likely represents the perturbation of immune targeting of self (black box). (C) Expanded view of dendrogram section in purple boxed region in (B), showing that transcripts cluster by functionality. Position of FKBP11 clustering with both interferon α and β receptor subunit 2 (IFNAR2) and interferon γ receptor 2 (IFNGR2).

To assess the propensity of the approach to report differences unrelated to sample group definitions, samples were randomly assigned into 2 groups of 16, each with 8 normal and 8 SARS-CoV-2 samples, and the number of transcript type-to-transcript type mean SM3DDs that passed FDR-control determined. Across 1000 reiterations of randomising samples, 3 reiterations reported that only 1 of 960,400 mean SM3DDs comparisons passed FDR-control, while the other 997 reiterations reported none.

To better visualize the results of the mean SM3DD comparison between SARS-CoV-2 and normal (Extended Data Table 1), we displayed the directional log10(P values) as a clustered heatmap, with ‘from’ and ‘to’ transcript type on alternate axes (Suppl. Fig. 2). The extent to which transcripts with shared function clustered together with very small cophenic distances was striking and most obvious for gene family members (Fig. 2C). Notably, FKBP11 clustered with both interferon α and β receptor subunit 2 (IFNAR2) and interferon γ receptor 2 (IFNGR2), while MZT2A clustered with the long noncoding RNA Metastasis Associated Lung Adenocarcinoma Transcript 1 (MALAT1), which is nuclear localised.

For prominent clusters of statistically shorter mean SM3DDs in SARS-CoV-2 samples, sets of clustered genes were assessed by Fisher’s Exact Test with FDR control for pathway level effects, utilising GeneCard Suite’s PathCard database ^12^. There are two bands where the mean SM3DDs are predominantly shorter for clusters of transcripts from which the SM3DDs were determined (FROM-Transcripts), indicating widespread effects. Pathway analysis identified that these likely represent ‘interferon γ signaling’ and extracellular matrix (ECM) deposition (SuperPathways ‘miRNA targets in ECM and membrane receptors’, ‘ECM proteoglycans’ and ‘extracellular matrix organization’) (Fig. 2B, Extended Data Tables 2-3). Assessment of transcript density for the five main induced collagen types confirmed extensive ECM deposition by pan-cytokeratin negative cells (Fig. 3A), while an assessment of the three interferon-induced transcripts in the assay confirmed widespread signaling (Fig. 3B).

**Figure 3.**
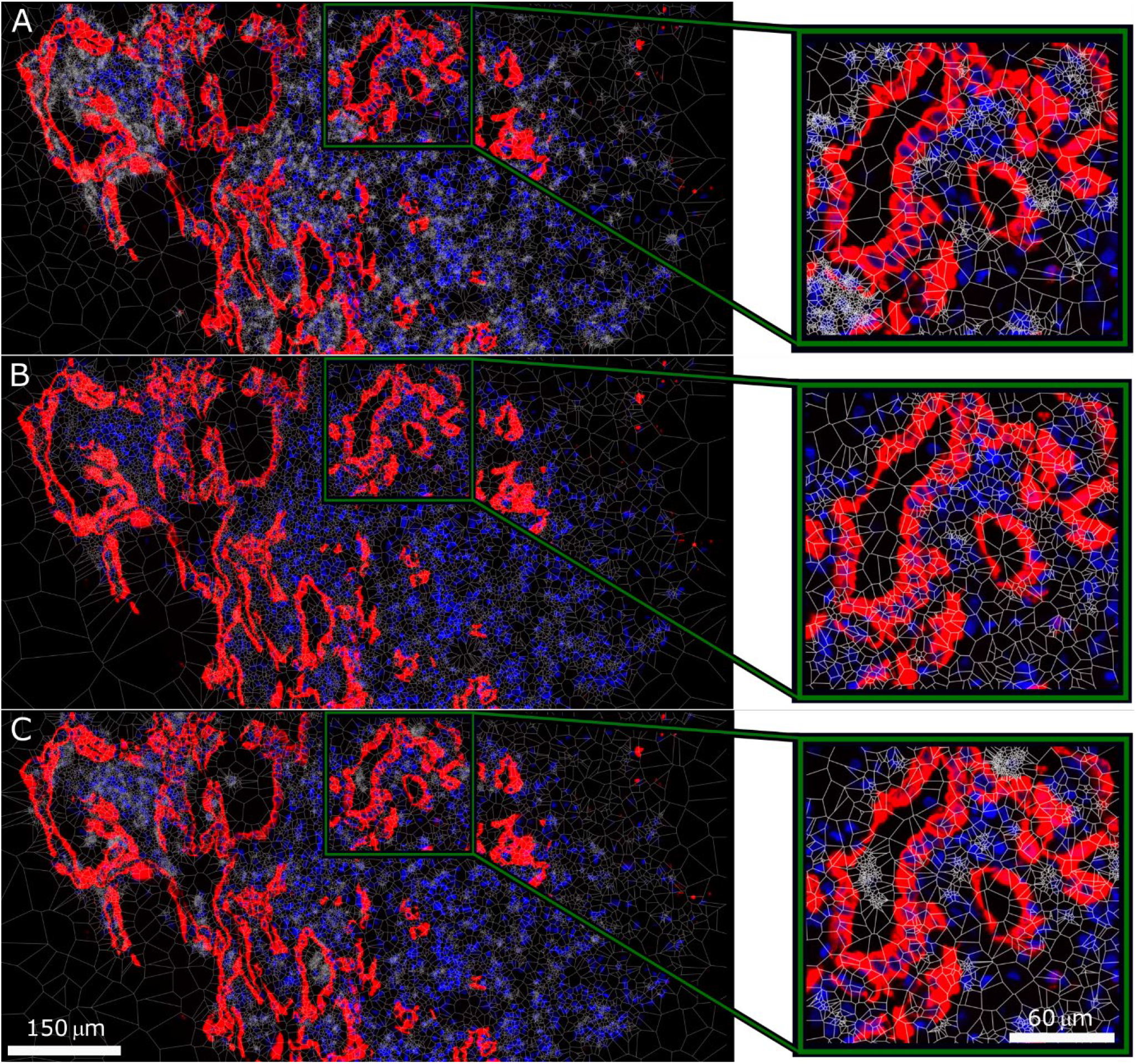
Transcript density across SARS-CoV-2 infected lung. Tessellation (white) of transcript density distribution for (a) COL1A1/COL1A2/COL3A1/COL6A2/COL6A3 showing ECM deposition by pan-cytokeratin negative cells, (b) IFI27/IFITM1/IFITM3 showing widespread interferon signalling, and (c) CD68/CD163/FCER1G/FCGR3A/GLUL/HLA-DPB1/HLA-DRB5/ITGB2/SPP1 showing a high density of cells involved in pathogen phagocytosis.

For one region of the clustered heatmap with statistically shorter mean SM3DDs in SARS-CoV-2 samples, pathway analysis reported an increase in microglia pathogen phagocytosis (‘phosphorylation of CD3 and TCR zeta chains’, ‘microglia pathogen phagocytosis pathway’ and ‘TCR signaling (REACTOME)’ for the cluster of FROM-Transcripts, and for the cluster of TO-Transcripts, ‘microglia pathogen phagocytosis pathway’ and ‘TYROBP causal network in microglia’) (Fig. 2B, Extended Data Tables 4-5). This result is limited though by the absence of any markers in the assay capable of distinguishing microglia from macrophages. Transcript density mapping for nine of the cluster of TO-Transcripts shows a high density of expressing cells, consistent with macrophage infiltration (Fig. 3C).

For one large cluster of TO-Transcripts, 107 SuperPathways passed FDR control. Notably, the third statistically strongest after ‘cellular responses to stimuli’ and ‘infectious disease’ was ‘SARS-CoV-2 infection’ (P = 5.44 x 10^-8^). Largely, the list of SuperPathways for this cluster likely represents a conglomerate of overlapping and interacting signaling pathways active or affected in SARS-CoV-2 infection (Fig. 2B, Extended Data Table 6).

For both another large cluster of TO-Transcripts and the cluster of FROM-Transcripts that specifically have shorter SM3DDs to them, no SuperPathways passed FDR control. Interestingly though, the cophenic distances across this cluster of FROM-Transcripts are notably shorter and functional similarities between several of the SuperPathways identified without FDR control suggest that this region of the heatmap may represent the perturbation of immune targeting of self (‘perturbations to host-cell autophagy, induced by SARS-CoV-2 proteins’ for FROM-Transcripts and ‘cancer immunotherapy by CTLA4 blockade’ / ‘FoxP3 in COVID-19’ for TO-Transcripts) (Fig. 2B, Extended Data Tables 7-8).

### Segmented PCA facilitates pathway identification

As any given transcript type may exist in multiple contexts within a tissue sample, we also implemented a PCA approach, segmenting the analysis of all SM3DDs according to the transcript type from which they were determined. For each of the 980 transcript-specific segments, the difference in means of the PCA scores between transcripts in SARS-CoV-2 and normal samples was determined within each PCA dimension. These transcript-specific differences were then grouped by PCA dimension and assessed for pathway level effects. Across each PCA dimension, the transcript-specific differences for genes in each PathCard SuperPathway were compared by t-test to those for all other genes. The FDR was again controlled by the two-stage Benjamini, Krieger, and Yekutieli procedure, both across each PCA dimension and, within each SuperPathway, across all PCA dimensions. This approach identified SuperPathways for the first four PCA dimensions, but not the six subsequent dimensions, as being affected by SARS-CoV-2 infection. This unsupervised approach identified SuperPathways self-evidently affected by SARS-CoV-2 infection, such as ‘infectious disease’, ‘innate immune system’, ‘interferon gamma signaling’ and ‘SARS-CoV-2 infection’, along with several related to ECM deposition, muscle contraction, fatty acid biosynthesis, nerve growth cone collapse and O_2_/CO_2_ exchange in red blood cells (Table 1). Mapping of muscle-specific transcripts to FOV images identified that ‘muscle contraction’ is likely a batch effect, probably caused by differences in sampling locations between the two groups (data not shown).

**Table 1.**
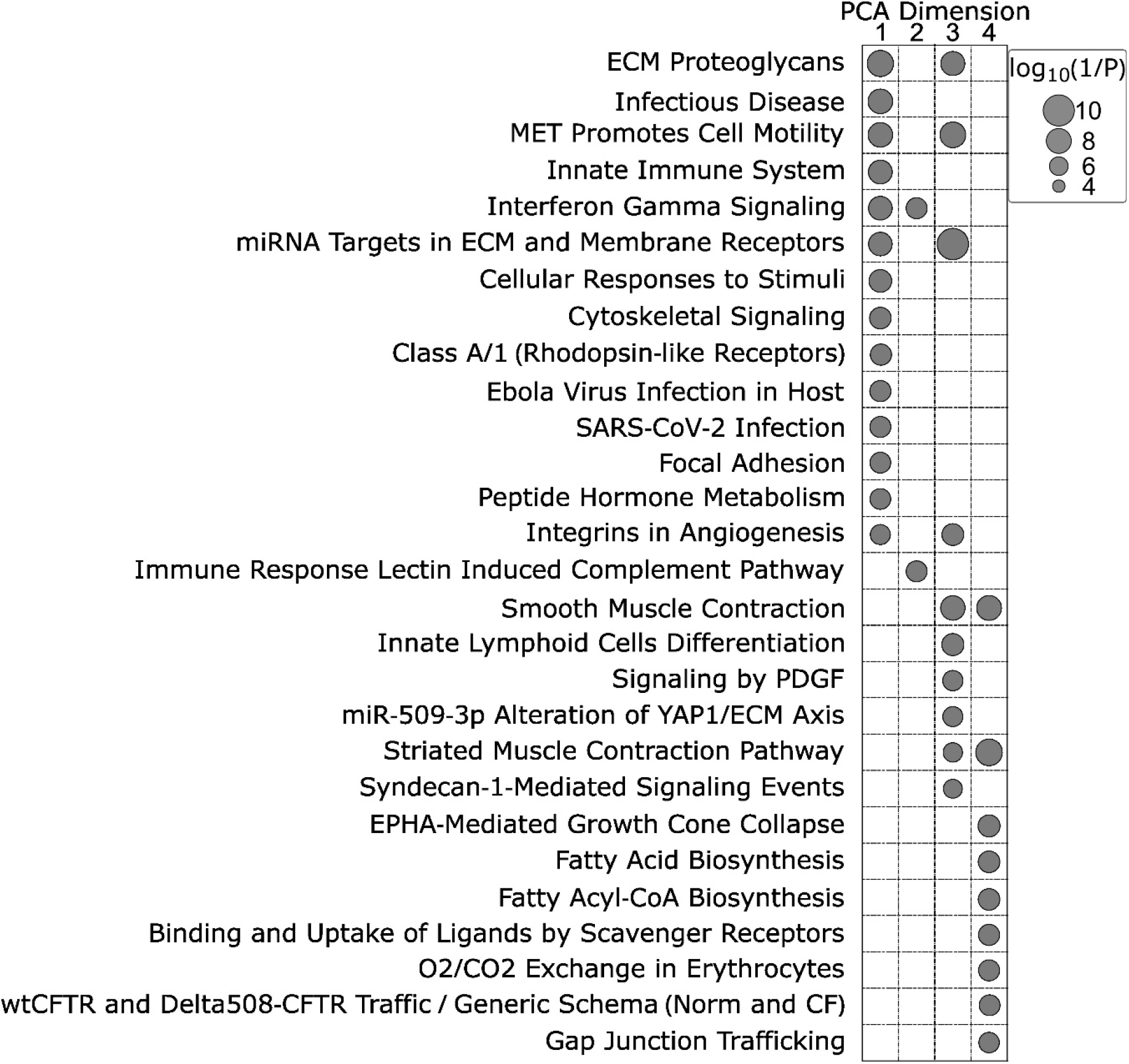
Sars-CoV-2 related SuperPathways identified by segmented PCA.

### Fibrotic Lung

To assess the versatility of SM3DD, we applied it to a lower-plex dataset with less divergent sample groups, choosing a publicly-available Xenium 343-plex spatial RNA dataset that included matched ‘more’ and ‘less’ fibrotic regions from 8 patients with pulmonary fibrosis ^8^. Clustering the results of a pair-wise mean SM3DD comparison (Extended Data Table 9) also lead to multiple instances of functionally-related transcripts clustering together despite having larger cophenic distances, such as AKR1C1 and AKR1C2; CD3D, CD3E and CD3G; CD8A and CD8B; HLA-DQA1 and HLA-DQB1; IFIT2 and IFIT3; IL1A and IL1B; KRT8 and KRT18; KRT15 and KRT17; and a proliferation-specific cluster CDK1, CCNB2, CENPF, TOP2A and MKI67. Pathway analysis of both the FROM- and TO-transcript clusters for the single prominent region of the heatmap with statistically shorter mean SM3DDs in ‘more’ fibrotic regions identified it as representing an unfolded protein response (Fig. 4, Suppl. Fig. 3, Extended Data Tables 10-11).

**Fig 4.**
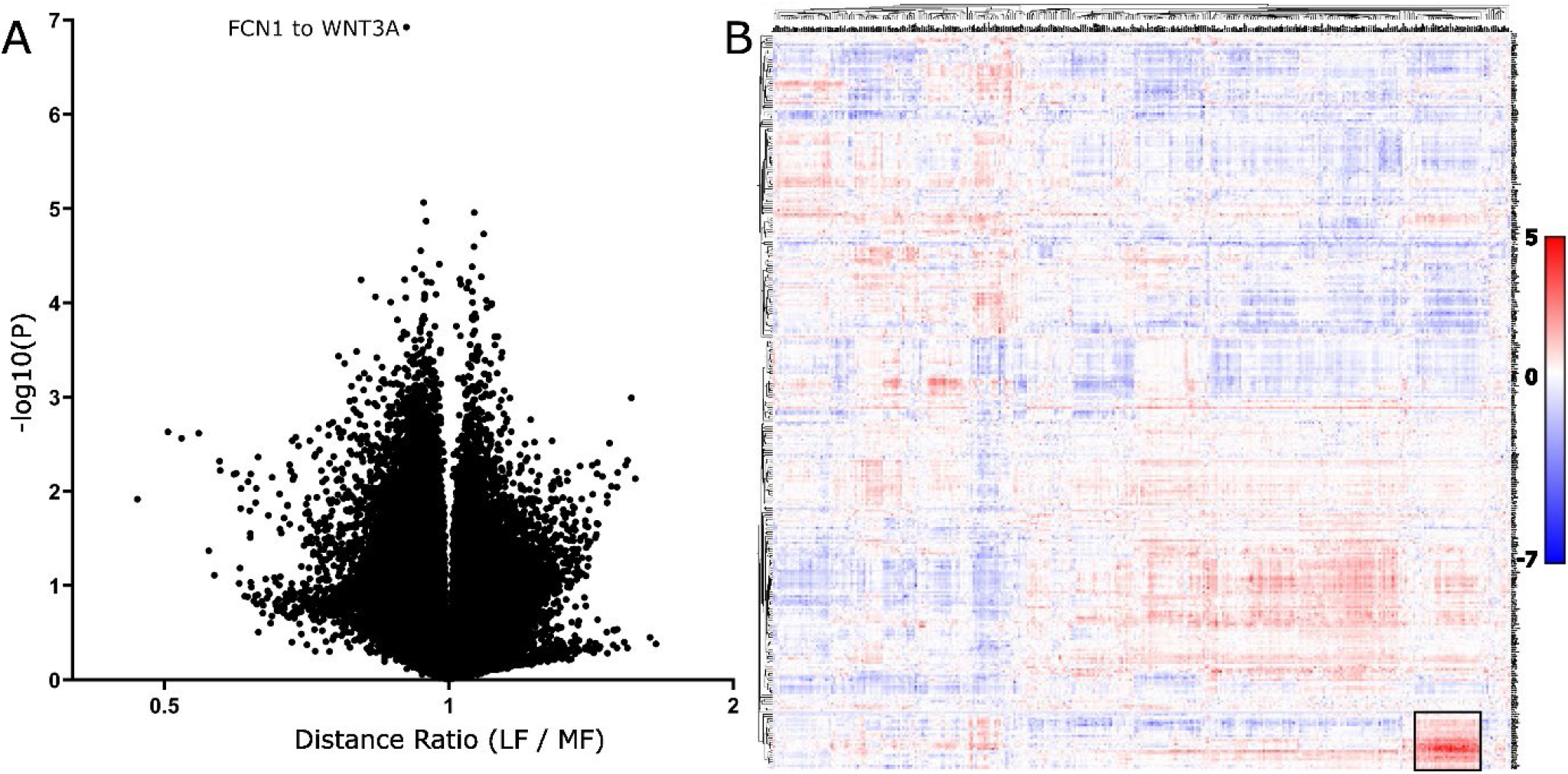
Comparison of transcript distances between ‘more’ and ‘less’ pulmonary fibrosis. (A) Volcano plot of all directional transcript distance comparisons. X-axis is distance ratio, where larger values represent shorter distances in regions that are more fibrotic. Y-axes are negative log10 (P values). (B) Clustering of directional negative log10(P values) from comparison of transcript distances between ‘more’ and ‘less’ pulmonary fibrosis. Scalebar indicates directional negative log10(P) value. The prominent cluster of transcripts with shorter distances (red) in more fibrotic regions were assessed by pathway analysis, identifying an unfolded protein response (black box).

Pathway analysis of transcript-specific differences in scores from the segmented PCA identified SuperPathways for the first two dimensions (Table 2). Four SuperPathways were identified for the first dimension, ‘defective CSF2RA causes {pulmonary surfactant metabolism dysfunction 4} SMDP4’ and the related ‘surfactant metabolism’, along with ‘FOXA1 transcription factor network’ and ‘prostaglandin synthesis and regulation’. Of the three SuperPathways identified for the second dimension, the strongest statistically was ‘triglyceride metabolism’ (P = 1.66 x 10^-14^), with the others both being related to it, ‘familial partial lipodystrophy’ and the ‘PPAR signaling pathway’.

**Table 2.**
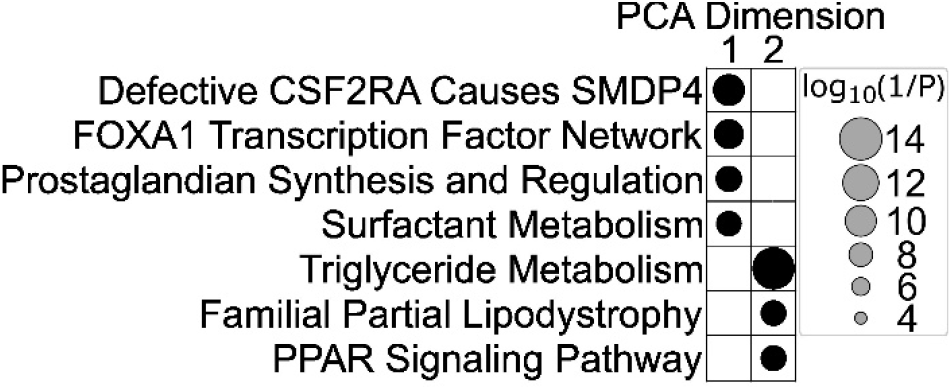
Pulmonary fibrosis related SuperPathways identified by segmented PCA.

## Discussion

A decrease in the mean SM3DD from FKBP11 to MZT2A transcripts was the most statistically significant difference between SARS-CoV-2 and normal samples. Clustering of directional NegLog10(P) values from mean SM3DDs comparisons grouped genes by functionality. FKBP11 clustered with interferon receptor subunits, implicating a potential role in SARS-CoV-2 infection for FKBP11 in either the facilitation or regulation of interferon signalling, which is consistent with previous findings in osteoblasts, where involvement of an FKBP11-CD81-FPRP complex in the expression of interferon-induced genes was reported ^13^. While not conclusive, a markedly shorter mean SM3DD from nuclear localised NEAT1-to-FKBP11, along with clustering of MZT2A with another nuclear localised lncRNA, MALAT1, alludes to the possibility that SM3DD may be detecting nuclear localised of FKBP11 mRNA during SARS-CoV-2 infection.

Pathway analysis of clusters coinciding with prominent regions of the heatmap exhibiting statistically shorter mean SM3DDs identified multiple biological processes affected by SARS-CoV-2 infection, with a high degree of consistency with previous studies. Widespread interferon γ signaling was identified by both the mean SM3DD and segmented PCA analyses and is known to be particularly associated with SARS-CoV-2-induced ARDS and fatal outcomes (reviewed in ^14^). Widespread ECM deposition was also identified by both analyses, consistent with other studies of autopsy samples from SARS-CoV-2 induced ARDS patients that reported extensive ECM remodelling with a prominent increase in collagen deposition ^15^.

While ‘microglia pathogen phagocytosis’ was identified, the assay does not include any microglia-specific markers assay, so this result may instead represent macrophage phagocytosis. Previous studies have reported the accumulation of inflammatory, profibrotic macrophages in the lung during SARS-CoV-2 infection (reviewed in ^14^) and in post-acute SARS-CoV-2 patients with persistent respiratory symptoms, the abundance of alveolar macrophages correlated with the severity of fibrosis, indicative of failed lung repair ^16^.

Albeit while not passing FDR control, a distinct region of the clustered results of the mean SM3DD comparison resulted in functional similarities between several SuperPathways related to the perturbation of immune targeting of self that included SARS-CoV-2-induced perturbation of autophagy and FoxP3 in SARS-CoV-2 infection. Multiple SARS-CoV-2 proteins are known to perturb host innate and adaptive immune systems (reviewed in ^17^). SARS-CoV-2 ORF3 specifically dysregulates autophagy ^18^ and pharmacological induction of autophagy reduced SARS-CoV-2 propagation in both primary human lung cells and intestinal organoids ^19^, while SARS-CoV-2’s S-protein dysregulates FoxP3-induced regulatory T cell development (reviewed in ^20^). The inclusion of ‘cancer immunotherapy by CTLA4 blockade’ in the associated SuperPathways alludes to possible overlap between viral and cancer cell induced immune perturbation.

While it has previously been reported that SARS-CoV-2 can infect and spread through the peripheral nerve system ^21^, this is the first report indicating that nerve growth cone collapse may be occurring in SARS-CoV-2 affected lung autopsy tissue. As neuropilin-1 can facilitate SARS-CoV-2 cell entry and infectivity ^22^ and has been shown to mediate semaphorin D/III-induced growth cone collapse ^23^, we speculate that nerve growth cone collapse may involve viral protein interaction with neuropilin-1.

Pathway level analysis indicated that ‘O_2_/CO_2_ exchange in red blood cells’ was affected by SARS-CoV-2 infection. While SM3DD does compute distances from hemoglobin transcripts in erythrocytes, the underlying differences found here are more likely to have been driven by changes in alveolar epithelial cells, which can be derived from hematopoietic stem cells and have been shown to express hemaglobin ^24^.

Pathway analysis of scores from the segmented PCA revealed fatty acid biosynthesis as being affected by SARS-CoV-2 infection. Lipids have been shown to accumulate in lungs and cells infected with SARS-CoV-2 and plasmid lipid pattern alterations correlate with disease progression and severity (reviewed in ^25^). Pharmacological inhibition of fatty acid synthesis for the treatment of SARS-CoV-2 infection was proposed following results where it blocked SARS-CoV-2 replication and improved survival in a mouse model ^26^.

The importance of surfactant homeostasis dysfunction in pulmonary fibrosis is well established ^27^, leading to the accumulation of floccular material in the alveoli and activation of an unfolded protein response ^28^, the latter of which was also identified here through pathway level analysis of the clustered mean SM3DD results. Pulmonary surfactant is a lipid/protein mixture that reduces surface tension at the air-liquid interface and is recycled by alveolar macrophages following granulocyte-macrophage colony-stimulating factor receptor (CSF2R) signalling ^29^. Pathway level analysis of segmented PCA score differences between matched ‘more’ and ‘less’ pulmonary fibrotic regions identified a change in the extent of surfactant metabolism dysfunction caused by defective CSF2R signalling, which can be due to neutralizing autoantibodies to CSF2 ^30^ or hereditary mutations in either subunit ^31^.

Pulmonary surfactant is ∼90% lipid by weight ^29^. The segmented PCA approach described here identified triglyceride metabolism as being differentially affected between matched ‘more’ and ‘less’ pulmonary fibrotic regions, along with two well described regulators of lipid metabolism, FOXA1 and PPAR signalling ^32, 33^. This is consistent with recent studies that have shown a strong association between lipid metabolism and both the onset and progression of pulmonary fibrosis, through the promotion of apoptosis and the induction of both pro-fibrotic biomarkers and endoplasmic reticulum stress ^34^, the latter of which can lead to an unfolded protein response.

Prostaglandin synthesis and regulation was also identified as being differentially affected between matched ‘more’ and ‘less’ pulmonary fibrotic regions. This is concordant with previous studies have reported on the role of prostaglandins in limiting pathological features of lung fibrosis, such as TGFβ-induced myofibroblast differentiation, along with lung fibroblast migration, proliferation and collagen deposition ^35^.

At a cellular level the data provided by spatial transcriptomics is low density. SM3DD overcomes the limitations inherent in low density data by leveraging extreme numbers of transcript-to-transcript distances. Pathway analysis of SM3DD’s output identified ‘SARS-CoV-2 infection’ without the assay including any SARS-CoV-2 transcripts, underscoring the power of the approach. We have shown that a segmented PCA approach has sufficient similarity across the transcript-specific segments for the first few PCA dimensions to facilitate pathway level analysis. The overall approach can assess changes in cell types typically missed by cell segmentation methods that rely on nuclear detection, such as erythrocytes and peripheral nerves, demonstrating here that SM3DD can lead to descriptions of peripheral nerve cell pathology without requiring either nerve cell segmentation or annotation, highlighting SM3DD’s utility in spatial transcriptomics. The signalling pathway and metabolic changes determined here from SM3DD analysis of the pulmonary fibrosis dataset were not reported in the dataset’s original publication ^8^, underscoring the value of including SM3DD as a complementary analysis. The power of SM3DD to discover new biology, from either inter- or intra-group comparisons, will likely grow with the advent of whole-transcriptome spatial assays ^36^.

## Extended Data Table and Supplementary Figure Legends

**Extended Data Table 1 Mean SM3DD comparison of SARS-Cov-2 vs normal lung**.

List of P values, directional negative log10(P) and ratios from t-test comparison of mean SM3DDs for SARS-CoV-2 and normal samples.

**Extended Data Table 2 SuperPathways indicating widespread interferon γ signaling**.

List of SuperPathways from GeneCard Suite’s PathCard database that passed FDR control for a prominent cluster of FROM-transcripts that had statistically shorter mean SM3DDs in SARS-CoV-2 samples, indicated by the dark green box in Figure 2B.

**Extended Data Table 3 SuperPathways indicating widespread extracellular matrix (ECM) deposition**.

List of SuperPathways from GeneCard Suite’s PathCard database that passed FDR control for a prominent cluster of FROM-transcripts that had statistically shorter mean SM3DDs in SARS-CoV-2 samples, indicated by the pink box in Figure 2B.

**Extended Data Table 4 SuperPathways indicating pathogen phagocytosis**

List of SuperPathways from GeneCard Suite’s PathCard database that passed FDR control for a prominent cluster of FROM-transcripts that had statistically shorter mean SM3DDs in SARS-CoV-2 samples, indicated by the light green box in Figure 2B.

**Extended Data Table 5 SuperPathways indicating pathogen phagocytosis**

List of SuperPathways from GeneCard Suite’s PathCard database that passed FDR control for a prominent cluster of TO-transcripts that were statistically shorter mean SM3DDs in SARS-CoV-2 samples, indicated by the light green box in Figure 2B.

**Extended Data Table 6 SuperPathways identifying SARS-CoV-2 infection and, predominantly, signaling pathways active during infection**.

List of SuperPathways from GeneCard Suite’s PathCard database that passed FDR control for a prominent cluster of TO-transcripts that were statistically shorter mean SM3DDs in SARS-CoV-2 samples, indicated by the blue box in Figure 2B.

**Extended Data Table 7 SuperPathways indicating that cluster may represent the perturbation of immune targeting of self**.

List of SuperPathways from GeneCard Suite’s PathCard database, identified without FDR control, for a prominent cluster of TO-transcripts that were statistically shorter mean SM3DDs in SARS-CoV-2 samples, indicated by the black box in Figure 2B.

**Extended Data Table 8 SuperPathways indicating that cluster may represent the perturbation of immune targeting of self**.

List of SuperPathways from GeneCard Suite’s PathCard database, identified without FDR control, for a prominent cluster of FROM-transcripts that were statistically shorter mean SM3DDs in SARS-CoV-2 samples, indicated by the black box in Figure 2B.

**Extended Data Table 9 Mean SM3DD comparison between ‘more’ and ‘less’ pulmonary fibrosis**.

List of P values, directional negative log10(P) and ratios from t-test comparison of mean SM3DDs for ‘more’ and ‘less’ pulmonary fibrosis.

**Extended Data Table 10 SuperPathway ‘unfolded protein response’ in regions with more pulmonary fibrosis**.

List of SuperPathways from GeneCard Suite’s PathCard database, identified without FDR control, for a prominent cluster of TO-transcripts that were statistically shorter mean SM3DDs in ‘more’ fibrotic regions, indicated by the box in Figure 4.

**Extended Data Table 11 SuperPathway ‘unfolded protein response’ in regions with more pulmonary fibrosis**.

List of SuperPathways from GeneCard Suite’s PathCard database, identified without FDR control, for a prominent cluster of FROM-transcripts that were statistically shorter mean SM3DDs in ‘more’ fibrotic regions, indicated by the box in Figure 4.

**Supplementary Figure 1 Comparison of transcript distances specifically to MZT2A between normal and Sars-CoV-2 infected lungs**. Volcano plot of comparisons of transcript distances specifically to MZT2A. X-axis is distance ratio, where larger values represent shorter distances in Sars-CoV-2 infected lungs. Y-axis is negative log10 (P values).

**Supplementary Figure 2 Full resolution of unmarked up heatmap from Figure 2B**

Clustered directional negative log10(P values) from comparison of transcript distances between normal and Sars-CoV-2 infected lungs.

**Supplementary Figure 3 Full resolution of unmarked up heatmap from Figure 4**

Clustered directional negative log10(P values) from comparison of transcript distances between ‘more’ and ‘less’ pulmonary fibrosis.

## Data availability

The extended tables in this study have been deposited to Zenodo: 10.5281/zenodo.15226673

## Acknowledgements

This study was supported by the Wesley Research Institute, the Australian Academy of Sciences and the University of Queensland. The authors would like to thank A/Prof Kirsty Short for their feedback. Funding for FSFG’s Laboratory was partially provided by the Medical Research Future Fund, Australia (with the support of the Queensland Children’s Hospital Foundation, Microba Life Sciences, Richie’s Rainbow Foundation, Translational Research Institute and The University of Queensland, Australia) under award number 2019485; and funding from the Cooper Rice-Brading Foundation, Australia, The Tie Dye Project, Bricks & Smiles, The Kids Cancer Project, Australia, Tour de Cure, the PA Research Foundation, the National Breast Cancer Foundation (award number 2023/IIRS0063). The content is solely the responsibility of the authors and does not necessarily represent the official views of the organizations and funding agencies.

## Author Contributions

Concept: TB, FSGF, AK. Experimental: JM, CC, EEK, LP, YK, YL, AN, ML, PSFG, SN, APCM, CMS, LN. Analysis: TB, AK2, AK. Writing and critical review: all authors *AK= Arutha Kulasinghe AK2=Aaron Kilgallon

