## Supplementary figures and images for "SM3DD with Segmented PCA: A Comprehensive Method for Interpreting 3D Spatial Transcriptomics"

### Supplementary Figure 1

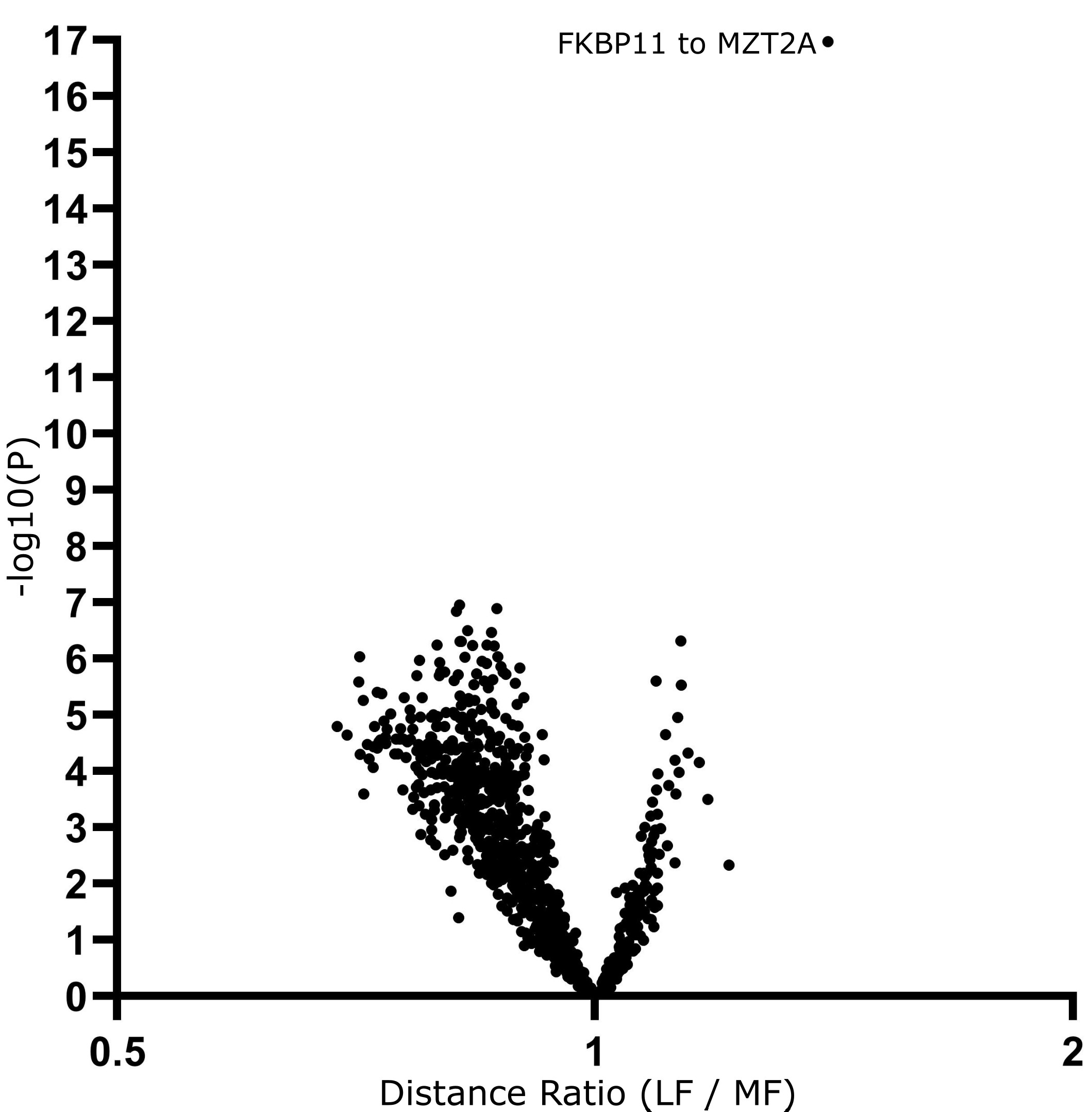

### Supplementary Figure 2

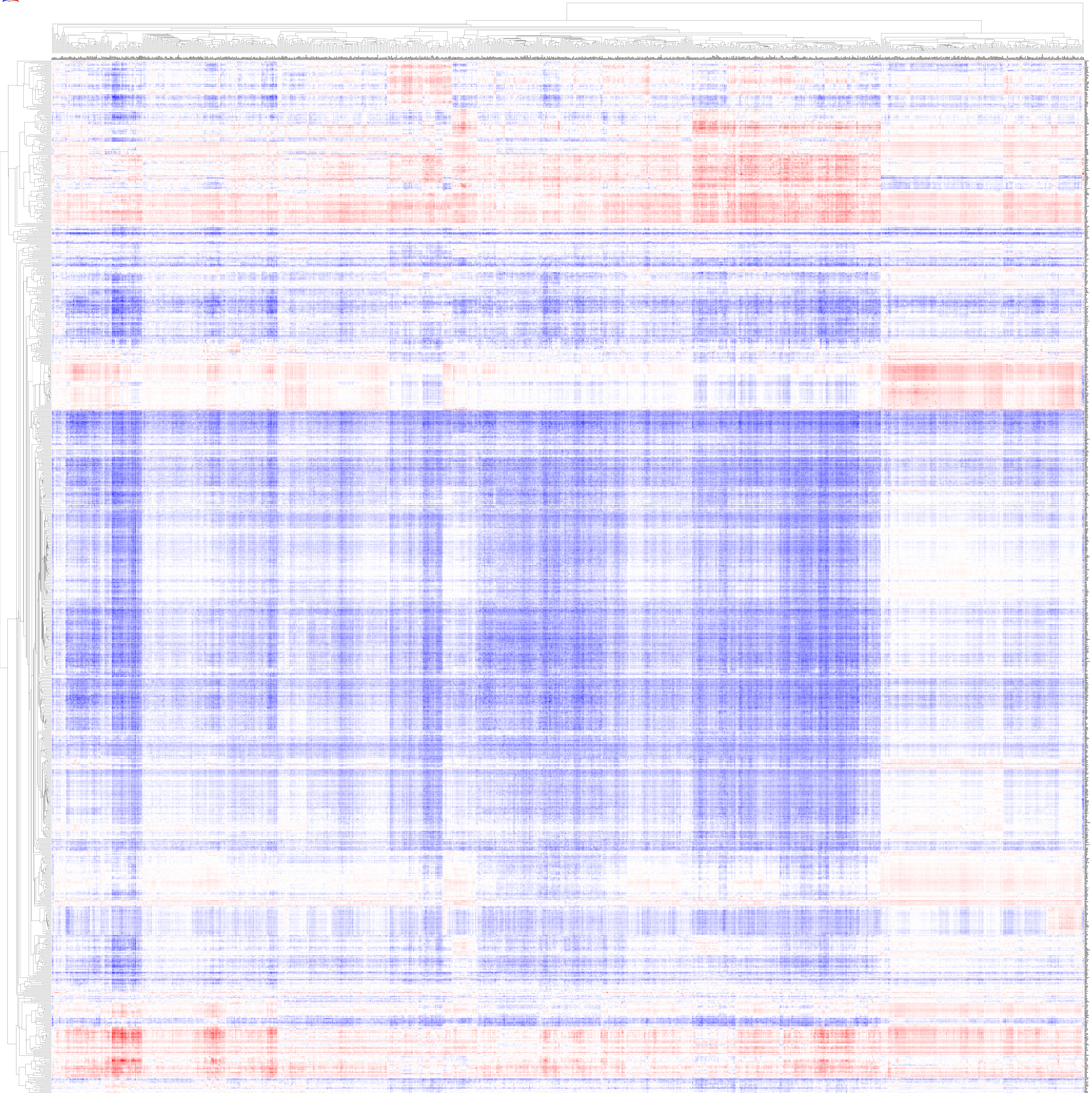

### Supplementary Figure 3

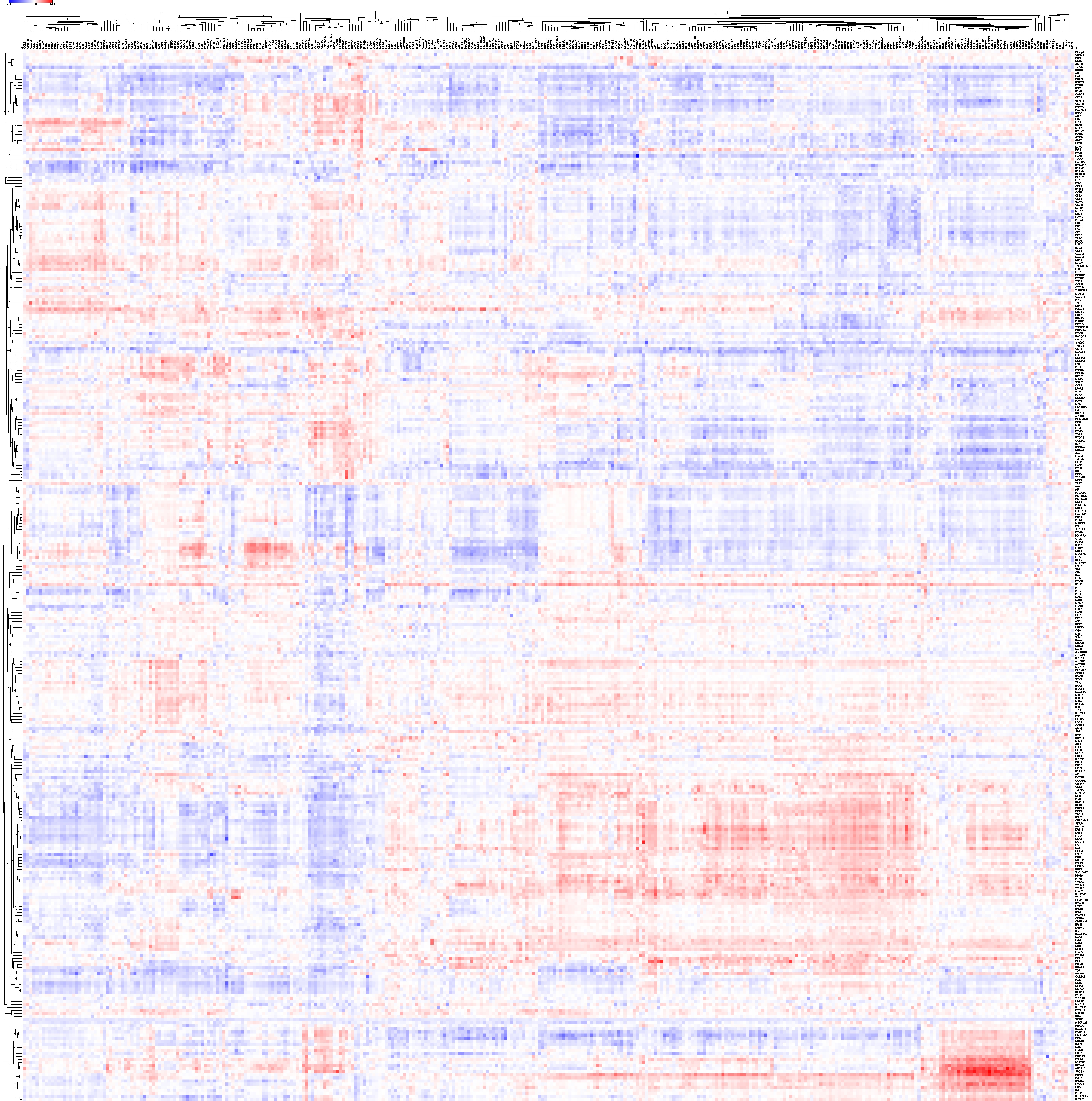
